# From microbial communities to regional biogeography: Unraveling patterns, determinants and the influence of bottom trawling in benthic microbiota

**DOI:** 10.1101/2023.08.09.552457

**Authors:** Guido Bonthond, Jan Beermann, Lars Gutow, Andreas Neumann, Francisco Rafael Barboza, Andrea Desiderato, Vera Fofonova, Stephanie Helber, Sahar Khodami, Casper Kraan, Hermann Neumann, Sven Rohde, Peter J. Schupp

## Abstract

Microbial composition and diversity in marine sediments are shaped by environmental, biological, and anthropogenic processes that operate on different scales. However, our understanding of benthic microbial biogeography remains limited. Here, we study how benthic microbiota vary at a regional scale in the North Sea with sediment characteristics, temperature, organic matter content, shear bed stress and bottom trawling intensity, a prevalent industrial fishing practice which heavily impacts benthic ecosystems. Using 16S rDNA amplicon sequencing, we characterized benthic microbiota from the top centimeter of 349 sediment samples and used uni-and multivariate statistical models, accounting for spatial autocorrelation, to disentangle the effects of the different predictors. Fitted models demonstrate how the geographic interplay of different environmental anthropogenic drivers shapes the structure and functioning of benthic microbial communities. Sediment properties were the primary determinants, with diversity increasing with sediment permeability but at the same time increasing with mud content, highlighting different underlying processes. Alpha diversity also increased nonlinearly with total organic matter content and temperature and showed a more complex relationship with bottom shear stress but decreased with bottom trawling intensity. These trawling associated diversity changes were accompanied by shifts in functional groups related to energy metabolism. Specifically, with increasing trawling intensity, we observed a transition toward more aerobic heterotrophic and less denitrifying metabolism. Our findings provide first insights of benthic microbial biogeographic patterns on a large spatial scale and illustrate how anthropogenic activity such as bottom trawling may influence the distribution and abundances of microbes and overall benthic metabolism at macroecological scales.

## INTRODUCTION

The biogeography of microbes is shaped by environmental, biological and anthropogenic processes that operate at different scales (Martiny *et al*. 2006). This includes the microbiota that colonize marine sediments in high cell densities (Musat *et al*. 2006; Probandt *et al*. 2018). Marine sediments filter and accumulate organic and inorganic matter and play a crucial role in the biogeochemical cycling of carbon, nitrogen, sulfur and metals (Middelburg 2018; Jørgensen *et al*. 2022). Most of these processes are carried out by microbes, which are organized at the microscale into communities that are tightly attached to sediment grains (Probandt *et al*. 2018). At the scale of millimeters to centimeters, microbiota are metabolically sorted along vertical redox gradients (Chen *et al*. 2017; Jørgensen *et al*. 2022). The aerobic heterotrophs that consume oxygen at the sediment surface are sequentially substituted underneath by anaerobes utilizing alternative electron acceptors, including nitrate, manganese/iron oxides, sulfate and finally carbon dioxide. Patterns in benthic microbial composition and diversity at macroecological scales (i.e., regional, continental and global; Gaston & Blackburn 2000), and how they arise from environmental drivers, have been studied less extensively (Shade *et al*. 2018) but correlate with sediment type (Probandt *et al*. 2017), temperature (Hicks *et al*. 2018), organic resource availability (Hoshino *et al*. 2020), primary production (Zinger *et al*. 2011), macrofaunal bioturbation (Laverock *et al*. 2014) and environmental disturbance (Galand *et al*. 2016).

While the benthic environment is subject to natural forms of disturbance driven by currents, waves and storms (van Denderen *et al*. 2015), it also experiences anthropogenic disturbances (Halpern *et al*. 2008), which potentially affect microbial biogeography. Bottom trawling, a prevalent fishing practice, represents the most extensive anthropogenic disturbance to seabed habitats (Kaiser *et al*. 2002). In the North Sea, over 60% of the overall bottom surface is trawled once or more per year (Eigaard *et al*. 2017). As large metal chains and heavy nets are dragged over the seafloor, the local environment is physically disturbed, resulting in a range of effects that act on different scales (Piet & Quirijns 2009). Trawling resuspends large amounts of sediment (Rijnsdorp *et al*. 2021; Breimann *et al*. 2022), alters seabed morphology (Puig *et al*. 2012), destroys biogenic structures (Tillin *et al*. 2006), and injures or kills benthic macrofauna (Bergman & Hup 1992; Bolam *et al*. 2014; Hiddink *et al*. 2017). Directly and indirectly, trawling may alter the benthic biogeochemistry by enhancing the oxygenation of buried organic matter, impacting the sequestration and remineralization of organic carbon (Falcão *et al*. 2003; Warnken *et al*. 2003; Morys *et al*. 2021; Sala *et al*. 2021; Epstein *et al*. 2022). It may also suppress benthic denitrification, and therewith potentially contribute to eutrophication (De Borger et al. 2020; Ferguson et al. 2020; Tiano et al. 2022). As bioturbation and bioirrigation activity by macrofauna controls benthic oxygen and carbon fluxes at regional scales (Neumann *et al*. 2021), bottom trawling may also affect benthic microbiota and microbial processes on larger temporal scales through its impact on macrofaunal populations, of which some species are highly susceptible to trawling (Reiss *et al*. 2009; van Denderen *et al*. 2015; Hiddink *et al*. 2017; Rijnsdorp *et al*. 2018).

Based on this range of trawling impacts on the benthic environment, benthic macrofauna and microbial processes linked with biogeochemical cycling, bottom trawling might also affect the composition and diversity of benthic microbiota at large scales. In this study, we present an analysis of regional scale patterns and putative determinants – including bottom trawling intensity – of benthic microbial diversity and composition in the central to southeastern North Sea. This marine region is known as one of the most heavily trawled regions in the world, but trawling intensities are highly heterogeneous (Eigaard et al. 2017; Kroodsma et al. 2018). We conducted 16S rRNA gene metabarcoding on 339 surface sediment samples from 149 sites across hundreds of kilometers, measured various environmental variables, obtained model-based estimates of bed shear stress levels (natural disturbance) and high resolution bottom trawling intensities recorded by the Vessel Monitoring System (VMS). To disentangle these variables, each representing different hypotheses, and evaluate the direction and shape of their relationship, we utilized uni-and multivariate statistical models, that control for spatial autocorrelation. Specifically, we were interested to evaluate the hypothetical effects of bottom trawling intensity on regional scale microbial biogeography.

## METHODS

### Sampling and sedimentological parameters

Sediment samples were taken with a 0.1 m^2^ van Veen grab (weight: 90 kg) during two scientific expeditions with the Research Vessel Heincke as part of an ecological long-term monitoring program (HE538, doi:10.2312/cr_he538 and HE562, doi:10.48433/cr_he562), in August 2019 and September 2020. In total, samples were collected from 150 stations in the Southeastern North Sea, including 50 stations on the Dogger Bank (HE538) and 100 stations in the Sylt Outer Reef (HE562, Figure 1). Sediment samples for DNA extraction were retrieved through a mesh lid on the top side of the grab to minimize disturbance and collected in 15 mL falcon tubes, taking the top centimeter only (HE538) or the top 1 to 10 centimeters as vertical core (HE562). In the latter case, three cores were taken approximately 10 cm apart in the same grab. Upon collection, tubes were stored at -20 °C on board, afterwards transferred to the laboratory in cooling boxes, and stored again at -20 °C. For granulometry, a subsequent sample was taken with a core (diameter: 4.5 cm, penetration depth approximately 6 cm) from the same grab. The bottom temperature was determined from a separate grab with a standard thermometer, inserted to approximately 3 cm below the surface. Forty grams (wet weight) of the sub-sample were dried, weighed and combusted for 5 hours at 500 °C. The total organic matter (TOM) content was calculated from the weight loss during combustion. The remaining sediment was fractionated in a sieve cascade with mesh sizes of 8000, 4000, 2000, 1000, 500, 250, 125 and 62.5 μm, corresponding to the Krumbein ϕ scale (Krumbein 1934). The mud (< 62.5 μm), sand (> 62.5 μm, < 2000 μm) and pebble (> 2000 μm) fractions were defined based on the percentage weight. For each sample, logistic regressions were fitted on the cumulative grain size distribution from which the median grain size was estimated by dividing the location and slope parameter multiplied by -1. The slope parameter was used as sorting coefficient.

**Figure 1.**
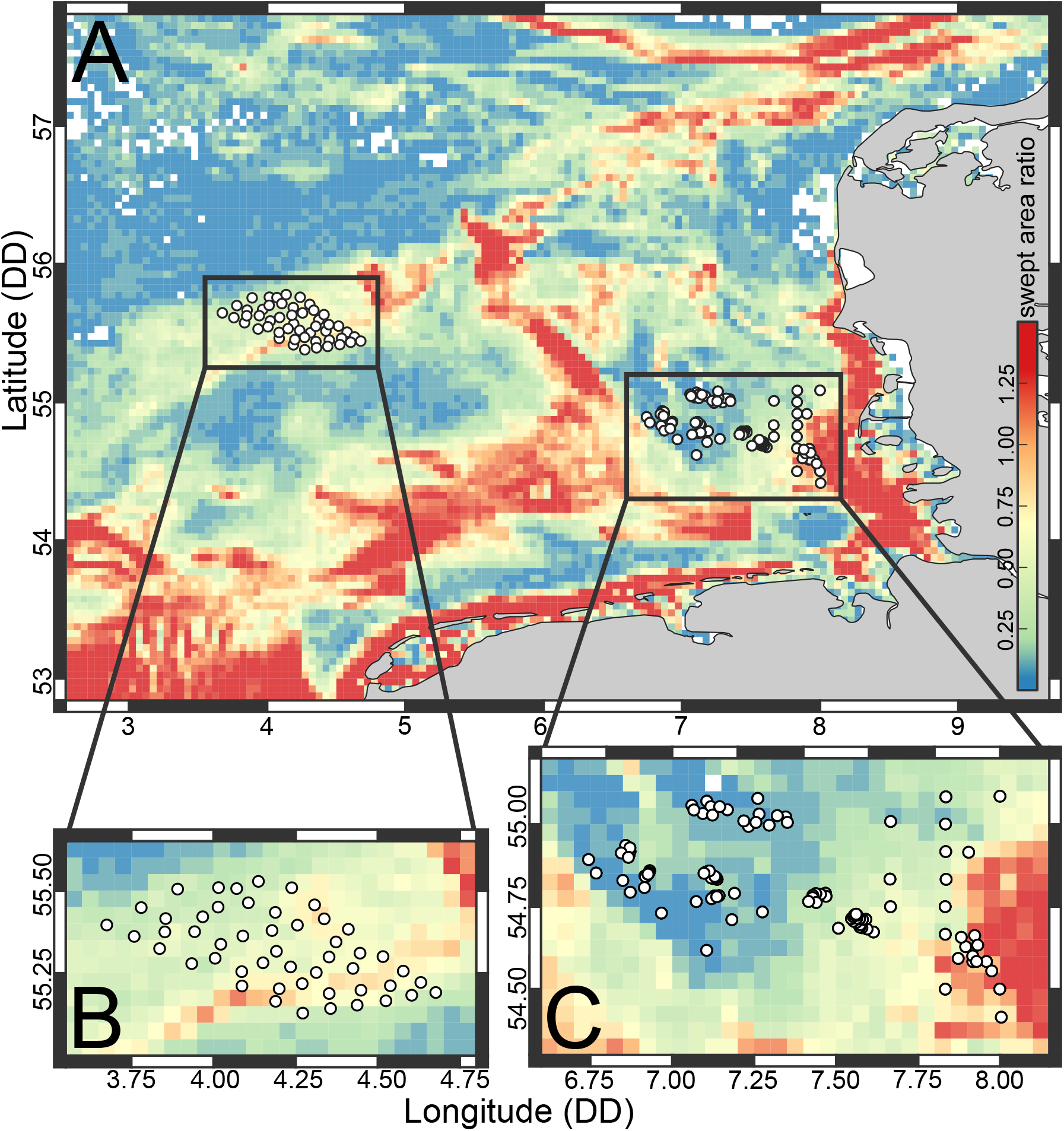
(A) Map of the central and south-eastern North Sea with stations sampled with van Veen grabs. The background heatmap displays the bottom trawling intensity calculated as swept area ratio (SAR) per year (see the main text for details on SAR). Panels B and C show zoomed sections of the two areas sampled on two different expeditions (HE538 and HE562).

### DNA extraction and library preparation

DNA was extracted from sub-samples of the top cm with the DNAeasy PowerSoil Pro kit (QIAGEN) following the manufacturer’s protocol. The 16S-V34 region was amplified with the primers 341F (S-D-Bact-0341-b-S-17) and 805R (S-D-Bact-0785-a-A-21; Klindworth *et al*. 2013), and two amplicon libraries were prepared following Gohl *et al*. (2016). The first PCR was run with the Phusion Green Hot Start II High-Fidelity PCR Master Mix (ThermoFisher) in volumes of 20 μL with 2 μL template and 0.5 μM of each primer, an initial denaturation step of 98 °C for 3:00 min, 20 cycles of 98 °C for 0:30 min, 55 °C for 0:30 min and 72 °C for 0:30 min and a final elongation step of 72 °C for 5:00 min. The second PCR was conducted in 20 μL volumes as qPCR with the SsoAdvanced Universal SYBR Green Supermix (Biorad) with the same program for 10 cycles. qPCR products were then pooled in equimolar volumes estimated from the C_q_ values, purified with a gel extraction step and sequenced on the Illumina MiSeq platform using the v3 600 cycle kit. Amplicon libraries included blanks from DNA extractions, negative PCR controls and mock communities (D6311, Zymo Research). FASTQ files were demultiplexed and further processed and quality filtered with MOTHUR (v1.46.1, Schloss *et al*. 2009), using the SILVA reference alignment (Quast *et al*. 2013, v132) for denoising and classification. Sequences were clustered into operational taxonomic units (OTUs) with the OPTICLUST algorithm based on the 97% similarity criterion. Mitochondrial, chloroplast, eukaryotic and unclassified OTUs, singleton OTUs, and samples with less than 1,000 reads were removed. The raw demultiplexed amplicon reads were deposited in the SRA database (accession: PRJNA988469). Metagenomes with KEGG Ortholog (KO) annotations (Kanehisa *et al*. 2014) were predicted with PICRUST2 v2.5.0 (Douglas *et al*. 2020).

### Statistical analysis

Eight variables were considered as potential drivers. As an estimate for contemporary bottom trawling intensity in the study area, we used subsurface (≥2 cm penetration of gear in the surface) fishing intensity from the OSPAR data & information management system (OSPAR 2017) which is based on data from the vessel monitoring system (VMS). The data are expressed as swept area ratio (SAR) at a resolution of 0.05 squared decimal degrees, averaged over the years 2009-2017. SAR-values indicate the number of times an area is fished with bottom-touching gears per time-period. As a measure of natural disturbance, we used bottom shear stress (in N/m^2^). Spatial data on shear stress (maximum value) were derived from the barotropic FESOM-C setup with tidal forcing for the North Sea. FESOM-C is the coastal sub-unit of the global Finite-volumE Sea ice Ocean Model (Androsov *et al*. 2019) and has been tested and verified in numerous idealized and realistic experiments for the North Sea (Fofonova *et al*. 2019, 2021; Kuznetsov *et al*. 2020; Sprong *et al*. 2020). The grid resolution varies from 30-100 m in nearshore areas to 1 km in deeper offshore areas. Tidal forcing was taken from the TPXO9 model (Egbert & Erofeeva 2002). The bottom friction coefficient varied spatially from 0.0025 to 0.003. The reference density field was averaged over the years 2019-2022 using salinity and potential bottom temperature for late August, which were taken from the NEMO setup for the North-West European shelf with a spatial resolution of 1.5 km (Copernicus Marine Service 2018). Further, temperature (°C), pebble content (% weight), sand content (% weight), mud content (% weight), median grain size (mm) and TOM (proportion), were used as predictors. These variables were measured as described above. Pebble, sand and mud contents were square root transformed, TOM content was logit transformed and the median grain size was transformed to the Krumbein ϕ scale, by taking the negative *log*_2_ of the particle diameter in mm. Correlations among these potential predictors were assessed based on Spearman’s rank coefficients (Figure S1). In cases of strong correlations (|ρ| > 0.7) only one of the correlated variables was included in the modeling process (Dormann *et al*. 2013).

Multivariate analyses were conducted with PERMANOVA (Anderson 2001) with the function *adonis2* from the R package VEGAN v2.6-4 and multivariate generalized linear models (mGLMs), assuming a negative binomial error distribution, using the *manyglm* function from the R package MVABUND v4.2.1 (Wang *et al*. 2012). To account for spatial autocorrelation, we followed the approach from Pelinson *et al*. (2021) and computed Moran eigenvector maps (MEMs) with the R package ADESPATIAL v0.3-20. The positive MEMs were included as artificial spatial variables in PERMANOVAs and mGLMs. In addition, the PERMANOVAs and mGLMs included the log transformed sequencing depth (LSD) as covariate, to control for differences in total read counts among samples, and the predictors of interest (i.e., median grain size, mud content, TOM, temperature, bottom shear stress and trawling intensity). The marginal effect of each predictor on the overall OTU, genus and KO composition was tested with PERMANOVA, using Aitchison distances and 9999 permutations. To consider the dependency among samples from the same grab, permutations were restricted to stations, by including station identity as a blocking factor. Based on the coefficients and standard errors from the mGLMs, we counted the number of OTUs and KOs with relative abundances varying with each predictor. Coefficients were considered significant when the theoretical 95% confidence region was entirely below (negative response) or above zero (positive response).

Non-metric multidimensional scaling (nMDS) was conducted based on Aitchison distances, using the R package VEGAN v2.6-4. To exclude the effect of the sequencing depth prior to the scaling procedure, partial residuals from the mGLMs were used. Ordination vectors for the predictors of interest were computed with the *envfit* function from the same R package.

Alpha diversity was measured in effective numbers (Jost 2006) of OTUs, genera and predicted KOs. As a measure of local scale beta diversity, we calculated Aitchison distances among cores from the same van Veen grab. Finally, we analyzed specific functional groups related to energy metabolism, using the summed predicted ortholog counts from complete or partial KEGG pathway modules. This included the KOs for cytochrome c oxidase to represent aerobic respiration (M00155), nitrification (M00528), denitrification (M00529), dissimilatory nitrate reduction and dissimilatory nitrite reduction (M00530), thiosulfate sulfate oxidation (M00595), dissimilatory sulfate reduction (M00596), methanogenesis from CO_2_ (M00567) and methane oxidation (M00174). Details on the KOs included in the modules are provided in Table S1.

Univariate responses were analyzed with generalized additive mixed models (GAMMs) using penalized cubic regression splines with the *gamm* function from the R package MGCV v1.8-41 (Wood 2011). First a global model was fitted, including all predictors (retained after the correlation analysis) as smooths with a maximum number of 3 degrees of freedom for each term. Station identity was included as random intercept to represent non-independence among samples from the same grab and the LSD was used as a covariate in the model to correct for the sequencing depth. To account for spatial autocorrelation, the models contained an exponential covariance structure, using the geospatial coordinates. Within grabs, the latitudinal coordinates of the outer samples were adjusted with ± 10^−6^ latitudinal decimal degrees to approximate the 10 cm distance between samples within a grab. Then, we compared the global model with the simpler models generated from all possible combinations of fixed effects, but always including the random effect and covariance structure, and selected the best model based on the AIC_c_ criterion. The relative importance (RI) of retained predictors was calculated as the sum of AIC_cw_ of all models with a ΔAIC_c_ ≤4 (Burnham & Anderson 2004).

## RESULTS

After quality filtering the raw reads, the total dataset counted 95,306 OTUs across 339 samples from 149 stations. 7,550 KEGG orthologs (KOs) were predicted from this dataset. At the genus level, the *Woeseia* (Gammaproteobacteria) was the dominant taxon in most samples, followed by unclassified Sandaracinaceae (Deltaproteobacteria), unclassified Actinomarinales (Actinobacteria), *Sva0996-marine-group* (Microtrichaceae, Actinobacteria), *Rhodopirellula* and *Blastopirellula* (Planctomycetes). Of these taxa, *Sva0996-marine-group* and the unclassified Sandaracinaceae (Deltaproteobacteria) were dominant in muddy sediments, whereas *Woeseia* (Gammaproteobacteria) and *Blastopirellula* (Planctomycetes) dominated in samples with high median grain size (Figure 2A, Table S2). Communities from areas with no or low trawling intensity tended to be dominated by unclassified Gammaproteobacteria and *Lutimonas* (Bacteroidetes) whereas samples from heavily trawled areas tended to be dominated by *Woeseia* and unclassified Sandaracinaceae (Gamma- and Deltaproteobacteria, respectively; see Figure 2B).

**Figure 2.**
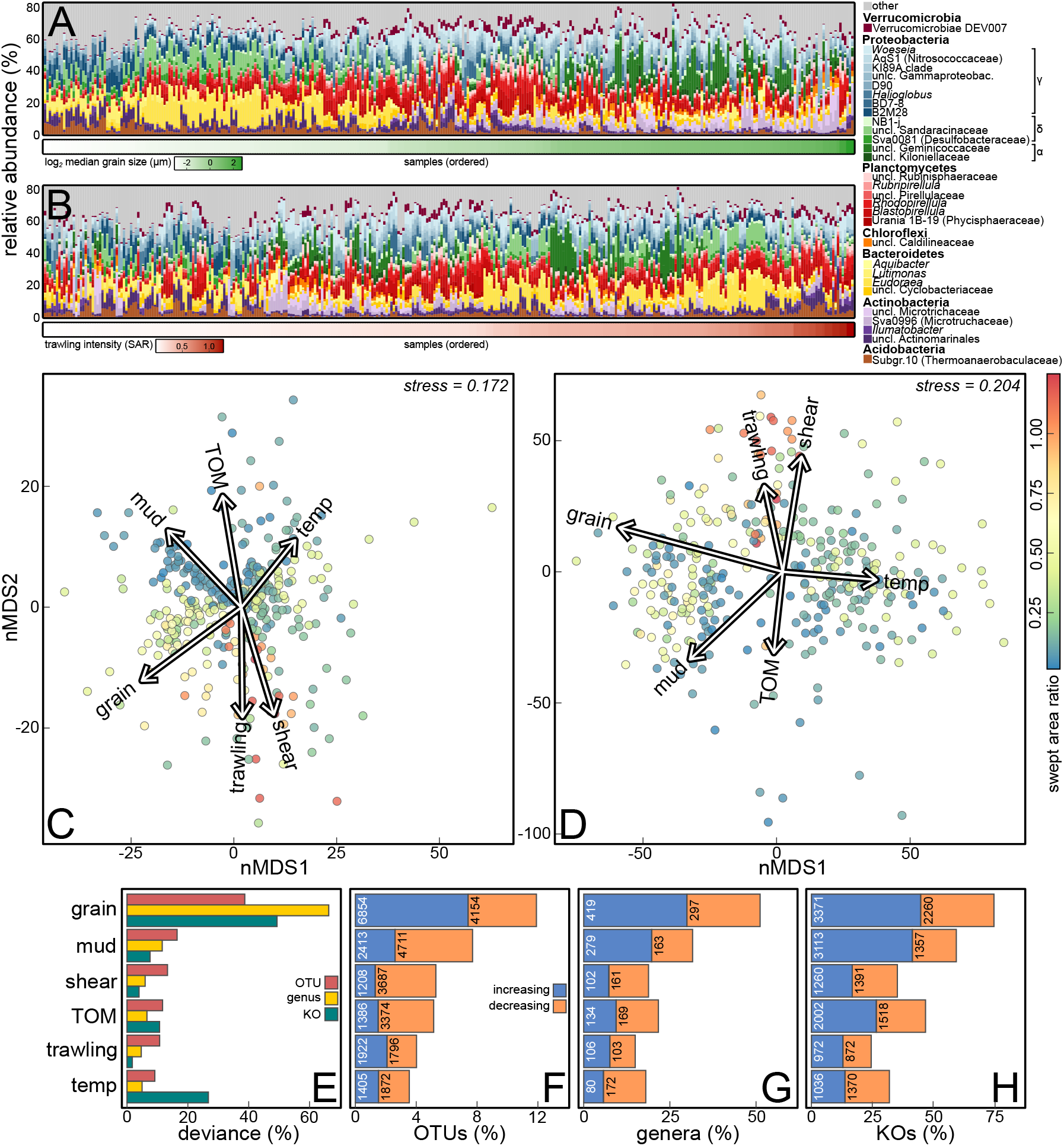
(A-B) Stacked bar plots of the 25 most abundant prokaryotic genera across all samples, (A) sorted by median grain size and (B) by swept area ratio. (C-D) nMDS plots based on Aitchison distances displaying compositional dissimilarities of the microbial communities after correcting for the sequencing depth, based on (C) OTUs and (D) KOs. The environmental variables were ordinated on the nMDS plots and are displayed as vectors. (E) The deviance explained by each term as percentage of the overall deviance explained by the model. (F-H) The number of (F) OTUs, (G) genera and (H) KOs, which differ in abundance with environmental predictors.

### Correlation analysis

Strong correlations (|*ρ*| > 0.7) were detected among pebble content, sand content, median grain size and sediment sorting. Mud content was correlated weakly with the other sediment variables (|*ρ*| < 0.5). Therefore, the model selection was conducted with the median grain size to represent the coarser sediment fractions and include mud content as separate variable. Moderate correlations were also detected between TOM content and shear stress (*ρ* = -0.62), TOM and trawling intensity (*ρ* = - 0.51), shear stress and mud content (*ρ* = -0.58) and shear stress and trawling intensity (*ρ* = 0.54). All other correlations were weak (|*ρ*| < 0.5, Figure S1).

### Community composition

Based on the geospatial coordinates, two positive MEMs were resolved, which were used as artificial spatial variables in the PERMANOVAs and mGLMs to account for spatial autocorrelation (Pelinson *et al*. 2021). Each of the included predictors had a significant marginal effect on OTU, genus and KO composition (p < 0.05, Table S3). These effects were also visible in nMDS plots (Figure 2C, D). For OTU, genus and KO composition, the median grain size was the most important predictor explaining 38.7%, 49.3% and 66.3%, respectively, of the overall explained deviance. Mud content was the next most important predictor for taxonomic composition (16.6% and 11.6%) and temperature for KO composition (26.7%). Bottom trawling intensity was less important but explained 10.7%, 4.7% and 1.7% of the overall deviance in OTU, genus and KO composition, respectively (Figure 2E). Similarly, the median grain size was also the predictor with which most OTUs, genera and KOs varied (11,008 OTUs, 716 genera, 5,631 KOs), followed by mud content. In total 3,738 OTUs, 209 genera and 1,844 KOs varied with bottom trawling intensity (Figure 2F-H).

### Diversity

For alpha diversity (effective OTU and genus numbers) the median grain size, mud content, temperature, bottom shear stress and trawling intensity were retained as informative predictors in the model selection (Figure 3A-J, Table S4-5). OTU diversity increased nonlinearly with the median grain size (Figure 3A, F) and mud content (Figure 3B, G). Both at the OTU and genus level, diversity increased with temperature (Figure 3C, H) and varied also with shear stress and bottom trawling. OTU and genus diversity decreased with bed shear stress from ∼ 200 to 300 N/m^2^ and then increased (Figure 3D, I). In response to trawling, OTU diversity dropped rapidly at relatively low trawling intensities (SAR < 0.25) but remained constant thereafter (Figure 3E). Genus diversity decreased linearly with bottom trawling (Figure 3J). For functional diversity (i.e., the effective numbers of predicted KOs), the model selection procedure yielded median grain size, mud content, temperature, TOM content and trawling intensity as informative predictors. Functional diversity increased linearly with median grain size, mud content and temperature (Figure 3K-M). In contrast to taxonomic diversity, the TOM content was retained and yielded a unimodal trend. Functional diversity decreased with trawling intensity (Figure 3O).

**Figure 3.**
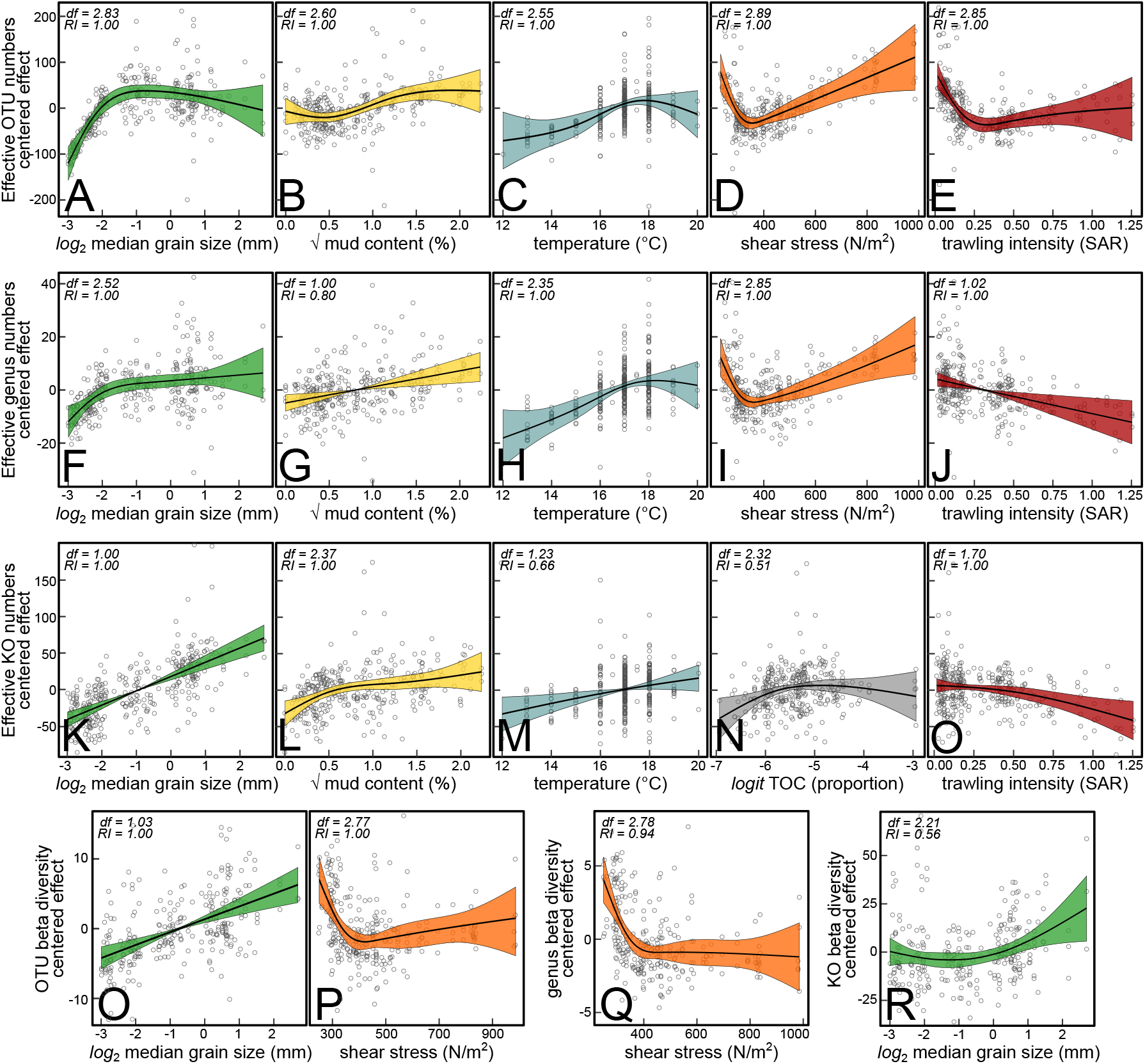
Partial effects estimated by generalized additive mixed models (GAMM) fitted on (A-E) OTU alpha diversity, (F-J) genus level alpha diversity, (K-O) KO alpha diversity, (O-P) OTU beta diversity and (Q) genus level beta diversity and (R) KO beta diversity, in response to the environmental predictors retained after model selection (see main text for further details). The vertical axes are centered and expressed in the scale of the response. The estimated degrees of freedom (df) of each smoot and the relative importance (RI) of each term are indicated in the top left corner. Shaded areas indicate the 95% confidence regions.

OTU beta diversity was best explained by median grain size and bottom shear stress, showing an increasing trend with median grain size and a rapid decrease followed by a gradual increase with shear stress (Figure 3O-P). At the genus level, only bottom shear stress was retained as informative predictor and reflected a rapid decrease at low stress levels (Figure 3Q). Median grain size was the only informative predictor for functional beta diversity, which increased with median grain size (Figure 3R).

### Predicted metabolic groups

Median grain size was retained in all models for metabolic groups except for aerobic respiration and dissimilatory nitrite reduction (Figure 4A). Mud content was an informative predictor for dissimilatory nitrite reduction, temperature for thiosulfate oxidation and dissimilatory nitrate reduction, and TOM content for predicted aerobic respiration, denitrification, dissimilatory nitrite reduction, dissimilatory sulfate reduction, thiosulfate oxidation and CO_2_ reduction. Further, bottom shear stress was retained for aerobic respiration, nitrification, dissimilatory sulfate reduction and methane oxidation and bottom trawling intensity for aerobic respiration (Figure 4B), nitrification (Figure 4C) and methane oxidation, denitrification (Figure 4D), dissimilatory nitrate reduction, sulfate reduction (Figure 4B) and thiosulfate oxidation.

**Figure 4.**
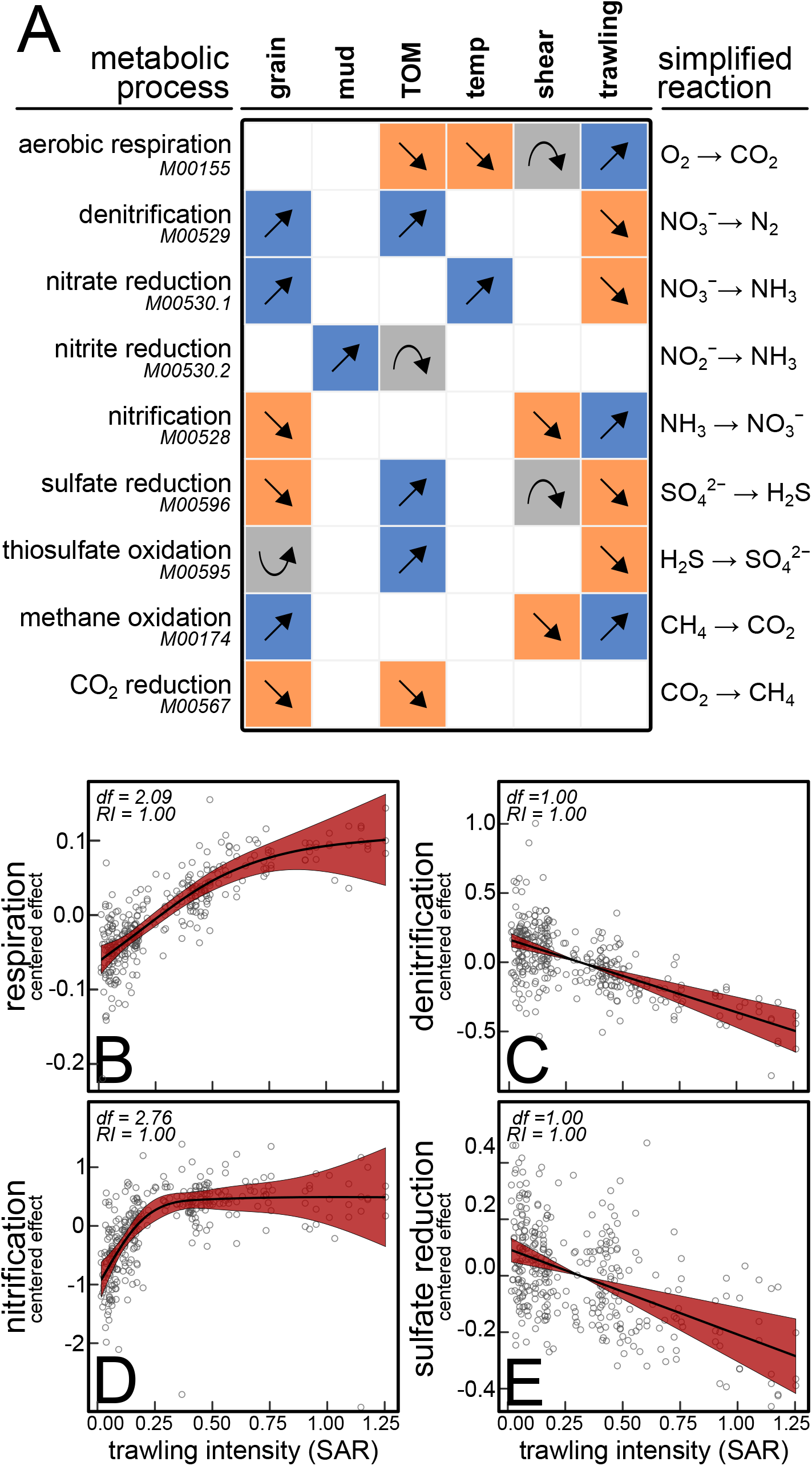
(A) Simplified trends estimated by GAMMs fitted on several predicted metabolic groups, based on KEGG gene modules (see main text for details). (B-E) Partial effect estimated by GAMMs for bottom trawling only on (B) predicted aerobic respiration, (C) nitrification, (D) denitrification and (E) sulfate reduction. The vertical axes are centered and expressed on the scale of the response (which was expressed as the natural logarithm of the sum of KO in the module). The estimated degrees of freedom (df) of each smooth and the relative importance (RI) of each term are indicated in the top left corner. Shaded areas show the 95% confidence regions.

## DISCUSSION

We characterized the microbial biogeography of the top sediment layer at a regional scale (i.e., across hundreds of kilometers) in the central to southeastern North Sea. All environmental variables included in our analyses partially explained at least some of the taxonomic and/or functional responses in benthic microbiota. This is in line with previous studies, which identified sediment characteristics, bottom temperature, and organic matter content as important variables shaping microbiota at larger scales (Wang *et al*. 2014; Probandt *et al*. 2017, 2018; Hoshino *et al*. 2020). Microbial communities were spatially autocorrelated, indicating the effect of unmeasured variables and/or historical contingencies. After correcting for environmental effects and spatial autocorrelation, we also found patterns in composition, diversity and functions associated with bed shear stress and bottom trawling intensity. Therewith, our work adds to previous studies which showed that bottom trawling impacts microbial biogeochemical processes (van de Velde *et al*. 2018; Tiano *et al*. 2019, 2022; De Borger *et al*. 2020) providing further evidence that bottom trawling may influence benthic microbial biogeography and metabolism.

### Granular properties

Sediment properties, represented by median grain size and mud content, were the primary determinant of benthic microbial diversity and composition. The median grain size and mud content explained different parts of the overall variation, indicating that these variables mirror different underlying processes possibly acting on different scales. The median grain size is known as a good proxy for permeability (Gangi 1985; Ahmerkamp *et al*. 2017), and therewith of porewater advection rates across sediments layers (i.e., centimeters), whereas mud particles may constrain advection at much smaller scales (i.e. micrometers) by forming cohesive aggregates wherein diffusion is the dominant process for solute transport. Mud aggregates could promote whole sediment diversity through the formation of microenvironments with functionally different communities, a process which is also known to occur in terrestrial soils (Bach *et al*. 2018). On the other hand, advection may promote alpha diversity rather through ecological processes such as dispersal and community coalescence between sediment layers and the water column. This may not only explain the observed patterns in alpha diversity, but also in beta diversity, for which only the median grain size but not mud content was a good predictor. Whereas the occurrence of mud aggregates likely has a specific effect occurring at microscales, advection also enhances the stochastic occurrence of microbes from the water column or deeper sediment layers, which may likely increase within-grab beta diversity. Others have reported a negative correlation between permeability and microbial diversity but proposed this was best explained by the typically higher organic content in muddy sediments (Franco *et al*. 2007; Probandt *et al*. 2017). Our analyses disentangle the effects of median grain size, mud content and TOM content by estimating partial effects of each predictor (i.e., how diversity varies with median grain size when levels of mud and TOM content are kept constant). Based on this, we suggest that the correlations reported in Franco *et al*. (2007) and Probandt *et al*. (2017) are most likely explained by confounding mud content, TOM content or both, rather than by permeability itself.

### Total organic matter content

At the global scale, organic matter content explains microbial community composition in marine sediments (Hoshino *et al*. 2020) as well as microbial biomass and diversity in terrestrial soils (Bastida *et al*. 2021). Our study in the North Sea confirms that benthic microbial composition and predicted functional diversity vary with TOM content. The increasing to unimodal trend in functional diversity agrees with macroecological theory which predicts that increased resource availability may promote diversity up to an optimum, beyond which biological processes (e.g., competition) favor specific functional groups at the costs of others (Geyer & Barrett 2019).

We also observed a relative decrease in aerobic respiration at higher levels of TOM content, accompanied by a relative increase in several anaerobic groups (Figure 4). This may mirror a transition from organic substrate limitation to oxygen limitation as resource availability increases, which could facilitate the proliferation of anaerobic heterotrophs, utilizing alternative electron acceptors. In the North Sea, high TOM content typically occurs in areas with low levels of near-bed turbulence and low permeability (Neumann *et al*. 2021). Here, absolute levels of oxygen consumption are relatively high (Neumann *et al*. 2019). Under these combined conditions of high TOM content and low oxygen supply through decreased advection, oxygen may be especially depleted near the sediment surface or in microenvironments such as dead-end pores and grain crevices, which may further boost anaerobic metabolism in the surface layer.

### Bottom temperature

The influence of temperature on the microbial composition and diversity is well established (Sharp *et al*. 2014; Zhou *et al*. 2016; Delgado-Baquerizo & Eldridge 2019; Li *et al*. 2023) and our results corroborate that bottom temperature is an important predictor for benthic microbial composition and alpha diversity. Thermal stress is often linked to microbial beta diversity in holobionts (Zaneveld *et al*. 2017; Ahmed *et al*. 2019; Bonthond *et al*. 2023; Li *et al*. 2023) according to the Anna Karenina Principle, which predicts beta diversity to increase due to a shift from deterministic to more stochastic processes acting on communities (Zaneveld *et al*. 2017). In a recent study on marine sediments, temperature was also identified as major driver of local scale beta diversity (He *et al*. 2021). Within the thermal range of this study (12°C to 20°C), we could not confirm a temperature beta diversity relationship. While temperature was an important predictor especially for functional composition (Figure 2E), only a relative decrease in aerobic respiration and increase in dissimilatory nitrate reduction were observed for the groups related to energy metabolism. Dissimilatory nitrate reduction may thermodynamically be more favorable at higher temperatures (Canion *et al*. 2014; Lai *et al*. 2021), which could explain this specific response, possibly hinting at a more prevalent role of dissimilatory nitrate reduction within benthic microbiota at higher temperatures.

### Mechanical disturbance

How microbial diversity varies along gradients of physical disturbance has received limited attention, but some studies found that diversity declines with disturbance in soils (Kim *et al*. 2013) or follows a unimodal trend in coastal marine sediments in agreement with the intermediate disturbance hypothesis (Galand *et al*. 2016). Within the range of disturbance levels estimated for the present study, we found a rather opposite relationship, with alpha diversity decreasing steeply toward a minimum at lower levels of disturbance, but increasing gradually. In surface sediments, near-bottom turbulence enhances pore water advection rates, but also promotes other modes of transport such as sediment erosion and mobile bedforms (Ahmerkamp *et al*. 2017). For example, the frequent resuspension of the surface sediment by mobile bedforms is constantly mixing individual grains into new redox zones and may prevent the local microbial community of a particular grain to complete succession to specific redox conditions. In this way, high bed shear stress levels could increase alpha diversity by preventing communities to reach successional stages where diversity is restricted by strong competition. Notably, bed shear stress and the median grain size were the only informative predictors for local beta diversity, yielding opposite trends. This may seem counterintuitive to the idea that higher advection rates (i.e., higher median grain size) promote beta diversity by increasing random occurrence of microbes from the water column or deeper layers. However, while bed shear stress indeed increases porewater advection rates, it also mixes and resuspends the sediment itself, potentially reducing substrate heterogeneity existing at the scale of centimeters or more (e.g., from macrofaunal bioturbation), which would simultaneously decrease within-grab beta diversity, as our results indicate.

The effect of bottom trawling on macrofaunal communities may overlap with the effects of natural disturbance (van Denderen et al. 2015; Rijnsdorp et al. 2018). Based on our findings, this may partially apply to microbial composition, as bottom trawling and bed shear stress were ordinated in similar directions in the nMDS (Figure 2C, D). At low levels of shear stress and bottom trawling, alpha diversity yielded also comparable trends but not at higher trawling rates. In contrast to bed shear stress, which was only a good predictor for taxonomic diversity, bottom trawling was also associated with decreasing functional diversity and different predicted metabolic groups, which may indicate differential effects of the two disturbance forms, reaching down to the functional level. While the nature of disturbance may be similar in some respects (e.g., sediment resuspension and mixing of the upper sediment layers), bottom trawling has different effects on the seabed morphology (Puig et al. 2012), and is more lethal to certain benthic macrofaunal species (van Denderen et al. 2015; Rijnsdorp et al. 2018).

### Bottom trawling

VMS based SAR estimates of fishing intensity are aggregated in space and time. Therefore, SAR values linked to datapoints do not necessarily reflect actual trawling events, as a given station that has never been trawled, may lay in a heavily trawled grid cell (Amoroso *et al*. 2018). Moreover, SAR does not distinguish short-and long-term effects, nor effects operating within and outside of trawling tracks and thus represent rather a probability that a geospatial position is in some way impacted by trawling. Consequently, this limits the ability to detect trawling effects and poses the need for high spatial replication to evaluate bottom trawling effects. In spite of this, benthic macrofaunal diversity and biomass have been found to decrease with SAR (Tillin *et al*. 2006; Hiddink *et al*. 2017; McLaverty *et al*. 2020). Here, taxonomic and functional microbial diversity followed similar trends in response to trawling intensities. The rapid decline in OTU diversity at limited trawling intensity (from 0 to 0.25 swept area ratio) may indicate a depletion effect, with the first trawling pressure having the highest impact and the effect of every subsequent trawl being proportional to the previous one (Pitcher *et al*. 2017). The more linear decrease in genus level and functional diversity suggests that the relatively high loss in OTU diversity at low trawling rates primarily concerns closely related and functionally redundant OTUs within a only few groups, while diversity at higher ranks responded more slowly.

The relative increase in predicted aerobic energy metabolism (i.e., aerobic respiration, nitrification) and decrease in anaerobic metabolic groups (i.e., denitrification, dissimilatory sulfate reduction), implies a trawling driven shift from anaerobic to aerobic heterotrophy within microbial communities. Microbial heterotrophy plays an important role in the storage and remineralization of organic carbon in marine sediments (Middelburg 2018; Jørgensen *et al*. 2022). Trawling may impede carbon storage or induce underwater carbon dioxide emissions by increasing aerobic microbial respiration (Sala *et al*. 2021), although in which quantities this occurs is currently subject of debate (Hiddink *et al*. 2023). Aerobic heterotrophy can be enhanced in different ways, including degradation of seabed morphology (Puig *et al*. 2012; Eigaard *et al*. 2017), re-exposure of buried organic matter to aerobic conditions (van de Velde *et al*. 2018; Tiano *et al*. 2019; De Borger *et al*. 2021), or by impacting macrofauna (Laverock *et al*. 2014; Epstein *et al*. 2022). While the redistribution of labile organic matter from deeper layers may fuel aerobic microbial heterotrophy (van de Velde *et al*. 2018; Morys *et al*. 2021), the putative trawling impact on burrowing macrofauna could further enhance aerobic microbial respiration, as benthic invertebrates consume most of the oxygen in the top sediment layer (Jørgensen *et al*. 2022) and transport labile organic matter to deeper anaerobic layers, allowing organic matter to escape aerobic microbial decomposition (Laverock *et al*. 2014; Neumann *et al*. 2021; Epstein *et al*. 2022).

A concurrent relative increase in nitrification and decrease in denitrification, implies also a shift in metabolism related to nitrogen cycling to be associated with trawling. Also this biogeographic trend aligns with biogeochemical patterns detected in previous studies that evaluated bottom trawling impact and reported reduced denitrification and increased sediment nitrate concentrations (Tiano *et al*. 2019; Ferguson *et al*. 2020; De Borger *et al*. 2021). Denitrification, which plays a major role in the removal of bioavailable nitrogen from marine systems such as the North Sea (Seitzinger *et al*. 2006), is maximized within marine sediments by the three-dimensional complexity of the redox structure which is formed by burrowing macrofauna (Laverock *et al*. 2014; Ferguson *et al*. 2020). Thus, both the mechanical trawling impact and the putative effects on macrofaunal communities, could explain the shift in nitrogen metabolism within benthic microbiota.

While the negative effects of bottom trawling on benthic macrofaunal diversity and biomass are well established for various systems, including the North Sea (Tillin *et al*. 2006; Reiss *et al*. 2009; Hiddink *et al*. 2017; Rijnsdorp *et al*. 2018), the susceptibility of macrofaunal communities to trawling is area specific and likely varies at different spatial scales (van Denderen *et al*. 2015). Therefore, the effects on the macrofaunal communities are not simply a function of SAR values and our results emphasize the need for future study to evaluate how the biogeography of benthic microbiota and macrofauna covary in the context of bottom trawling.

## Conclusions

Our models successfully explained regional scale environmental patterns in benthic microbial biogeography. Generally, sediment characteristics were identified as the most important determinants of structure and functioning. We posit that both alpha and beta diversity are shaped by processes operating at different spatial scales, with mud content enhancing whole sediment diversity by the formation of microscale environments, permeability (represented by the median grain size) promoting alpha and beta diversity by increasing the exchange of microbial cells across sediment layers at the centimeter scale and bed shear stress operating at the scale of meters or more, reducing local scale substrate heterogeneity and beta diversity in parallel.

After accounting for environmental predictors, bed shear stress and spatial autocorrelation, microbiota varied also with bottom trawling, mirroring a decrease in alpha diversity with increasing trawling intensity. These patterns were partially similar to those of bed shear stress, potentially indicating overlapping effects of the two types of physical disturbance, but were associated with different functional changes. Specifically, two noteworthy patterns in energy metabolism emerged, including a relative shift towards more aerobic metabolism and from denitrification to nitrification. While our data cannot be translated to chemical fluxes, and may not necessarily indicate changes in biogeochemical cycling, they do reveal notable biogeographic trends at a regional scale, which have not been documented before. Finally, the results of this study emphasize that the microbial biogeographic consequences of bottom trawling activity at the global scale merit further research.

## DATA AVAILABILITY

The raw de-multiplexed V4-16S gene amplicon reads and associated metadata are available from the SRA database under the Bioproject accession number PRJNA988469.

## ACKNOWLEDGEMENTS

The authors are grateful to the crew of the research Vessel FS Heincke for their support during the expeditions in 2019 (HE538) and 2020 (HE562) and Birgit Brinkmann and Petra Schwarz for assistance in the laboratory at the ICBM at the University of Oldenburg.

## FUNDING

This study was funded by the German Federal Ministry of Education and research (BMBF), through the DAM pilot mission: MGF North Sea (Grant number 03F0847B). JB and LG were funded by the German Federal Agency for Nature Conservation (BfN; grant number 3519532201, LABEL project).

## CONFLICT OF INTEREST

The authors declare that they have no conflict of interest.

## SUPPLEMENTARY FILES

Table S1. KEGG pathways used in the analyses

Table S2. Dominant genera

Table S3. PERMANOVA tables

Table S4. AIC_c_ tables

Table S5. GAMM output

Figure S1. Correlation matrix based on spearman rank coefficients

## Notes

### Competing Interest Statement

The authors have declared no competing interest.

