## Supplementary figures and images for "From microbial communities to regional biogeography: Unraveling patterns, determinants and the influence of bottom trawling in benthic microbiota"

### Figure S1

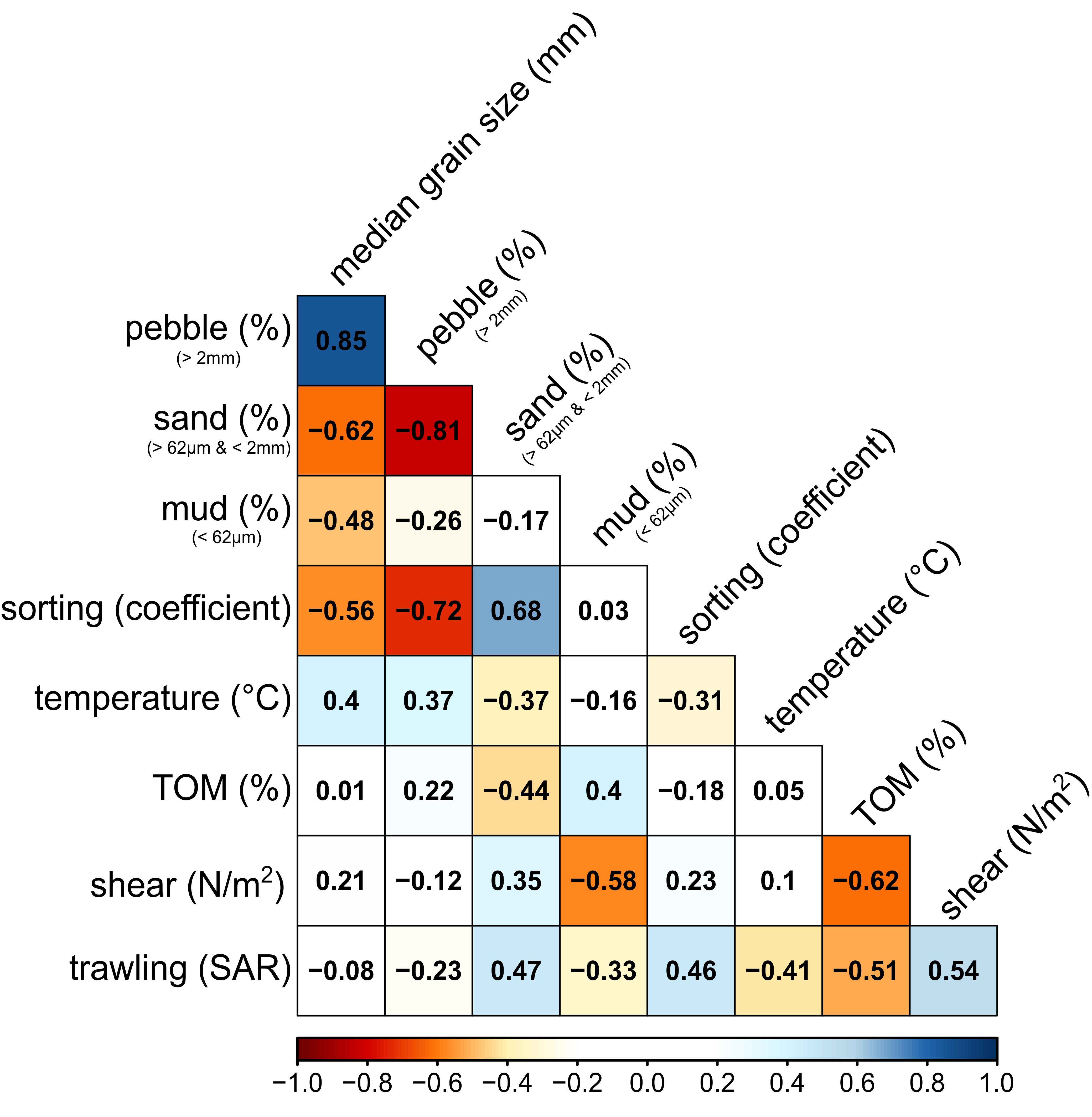
